# In vivo evaluation of thermally drawn biodegradable optical fibers as brain neural interfaces

**DOI:** 10.1101/2024.04.08.588517

**Authors:** Parinaz Abdollahian, Kunyang Sui, Guanghui Li, Jiachen Wang, Cuiling Zhang, Yazhou Wang, Rune W. Berg, Marcello Meneghetti, Christos Markos

## Abstract

Optical fiber technology has a critical role in modern neuroscience towards understanding the complex neuronal dynamics within the nervous system. In this study, we manufactured amorphous thermally drawn poly D, L-lactic acid (PDLLA) optical fibers in different diameters. These fibers were then implanted into the lateral posterior (LP) region of the mouse brain for 4 months, allowing us to assess their degradation characteristics. The gradual dissolution of the implanted PDLLA optical fibers in the brain was confirmed by optical microscope and scanning electron microscopy (SEM), molecular weight measurements, and light transmission spectroscopy. The results indicate that the degradation rate was mainly pronounced during the first week. Degradation after 4 months resulted in the formation of micropores on the surface of the implanted fiber within the gray matter region of the brain. Moreover, the current PDLLA optical fiber implant offers efficient light transmission in the short-wavelength near-infrared (SW-NIR) range. SW-NIR enables deeper tissue penetration and reduced light scattering, making it ideal for optogenetics and functional imaging with therapeutic potential in neurological disorders. We believe that the provided PDLLA optical fiber in this study constitutes a promising candidate for the development of next-generation biocompatible, soft, and biodegradable bi-directional neural interfaces

## Introduction

The use of light in brain research has revolutionized our understanding of the underlying mechanisms of neural circuitries and paved the way for innovative investigations, ranging from optogenetics ^1^ and imaging ^2^ to functional near-infrared spectroscopy (fNIRS) ^3^, photodynamic therapy ^4^, and light-based therapy ^5^. Optical brain implants play a crucial role in optogenetics, which is a powerful technique for interrogation and control of neural activity in millisecond timeframes ^6^. Typical optical implants are relying upon optical fibers or wave-guides that are carefully implanted into specific brain regions to deliver light to targeted neurons expressing light-sensitive proteins (opsins) ^7^. However, several design and material considerations should be addressed in the development of brain implants and neural interfaces. For instance, the implant must be not only bio-compatible but with low Young’s modulus to minimize tissue damage and inflammatory responses ^8^. This typically involves using materials as close as possible to the brain tissue, such as biocompatible polymers ^9^ or soft glasses ^10^. Also, neural interfaces should be flexible and minimally invasive to be inserted into brain tissue with minimal disruption. Thin, tapered optical fibers ^11^ or microscale waveguides are often used to target specific brain regions and deliver light precisely ^12^. Optical fibers or waveguides based on low-loss materials are used to maximize the amount of light reaching the targeted neurons ^13^. Polymer optical fibers (POFs) in particular have attracted significant attention for the development of neural interfaces because they can fulfill the aforementioned requirements but also can be tailored to accommodate several advanced materials with unique multi-functionality. POFs are typically based on standard thermoplastic polymers (polymethylmethacrylate, polycarbonate, cyclic olefin copolymer) and they have been extensively used for the development of fiber sensors and tunable devices ^14–16^ and more recently have attracted significant attention as fiber-based neural interfaces ^17–24^, However, an additional critical factor that has a role towards the development of the next-generation neural interfaces is their biodegradability, which can avoid an extra post-surgical step while suppressing the long-term inflammation response. As the biodegradable implant gradually breaks down over time, it can be absorbed by the surrounding tissue or metabolized by the body, minimizing the adverse immune reactions ^25^. Another advantage of biodegradable neural implants is their compatibility with tissue growth and regeneration. As the implant degrades, it creates space for natural tissue regeneration, enabling the integration of new tissue into the implant site. This property can be particularly beneficial in cases where the goal is to restore damaged neural tissue or create a bridge for neural regeneration ^26^.

Biodegradable polymers that find application in biomedical device technology can be categorized into two main groups including naturally occurring biopolymers such as gelatin ^27^, cellulose ^28^, and silk ^29^, and synthetic polymers including alginate polyacrylamide ^30^, polyethylene glycol (PEG) ^31^, polycaprolactone(PCL) ^32^, poly(acrylamide-co-poly(ethylene glycol) diacrylate) (p(AM-co-PEGDA)) ^33^, poly(lactic-co-glycolic acid (PLGA) ^34^, poly(L-lactic acid) (PLLA) ^35^, and PDLLA ^36^. Amongst them, polylactic acid (PLA) derivatives such as PLLA, PLGA, and PDLLA have gained significant attention in the field of biodegradable polymers for various biomedical applications ^37^. PLA derivatives are bio-compatible and biodegradable polymers derived from renewable sources like cornstarch or sugarcane ^38^, known to be well-tolerated by the human body and to not induce significant immune responses or adverse reactions when implanted ^39^. These polymers naturally degrade in the body through hydrolysis into non-toxic lactic acid, which can be metabolized. This property is advantageous for temporary optical fiber implants, as the body can safely resorb them without the need for surgical removal ^40^. Furthermore, the degradation rate and mechanical properties of PLA derivatives can be tailored by modifying their molecular weight or crystallinity. This allows for the customization of implants to match specific application requirements ^41^. Previously, researchers used PLA to develop microneedle arrays for enhanced percutaneous light delivery in dermatologic therapies, where light can be focused through the needle tips. Experiments with bovine tissues demonstrated a 9-fold increase in light delivery efficiency ^42^. However, this kind of microneedle device is not suitable for the stimulation of deeper structures, as those required for brain neuromodulation. PLGA has also been used to produce implants. Despite biocompatibility and biodegradability, however, these implants demonstrate early foreign-body reactions, occurring at 3 to 6 months following implantation, depending on the polymeric ratios.^43^ In 2018, Fu et al. fabricated PLLA biodegradable optical fibers for neural implants through a straightforward thermal drawing process, exhibiting flexibility and optical transparency. In vitro degradation studies show their structural evolution and optical property changes, while in vivo experiments demonstrate biocompatibility and full degradation. These PLLA fibers enable deep brain fluorescence sensing and optogenetic applications, showcasing their potential for fully biocompatible and bioresorbable POFs in biomedicine ^44^.

PLLA, however, exhibits a highly crystalline structure, causing its biodegradation to be slower than the one of amorphous PLA derivatives, whose random molecular arrangement makes them more susceptible to hydrolytic degradation ^43^.

Recently, Gierej et al. fabricated microstructured biodegradable and biocompatible POFs using amorphous PDLLA. They investigated the influence of polymer processing on PDLLA’s molar mass decrease and revealed varying degradation rates in vitro ^45^.

In this work, we developed a thermally drawn bio-degradable PDLLA POF. In order to enhance the degradation and mechanical properties, the starting PDLLA bulk was fabricated using a cast-quench technique, which we have shown to decrease the crystallinity of the material. The broad optical transmission of the fabricated fiber was experimentally evaluated, and its degradation was investigated both in vitro in phosphate-buffered saline solution (PBS) and in vivo in the brains of freely moving mice.

## Results and discussion

### Polymer casting and fiber drawing

The thermal history of PDLLA during its fabrication is a crucial factor in determining its degree of crystallinity ^29^. This is an important parameter to keep into account when fabricating implantable biodegradable devices, since a lower degree of crystallinity results in both an increase in degradation rate and a decrease in Young’s modulus, which translates in a smaller foreign body response from tissue ^30 31^. Furthermore, amorphous polymers offer numerous advantages over crystalline polymers for optical fiber fabrication: they are easier to process, ensuring uniformity and consistency during fiber production ^46^, and highly transparent thanks to the reduced scattering losses ^47^. Moreover, the flexibility and elasticity of amorphous polymers facilitate easier handling and installation of optical fibers, reducing the risk of breakage and mechanical stresses ^48^.

A well-known method to produce amorphous materials from melts is thermal quenching, where a rapid cooling process prevents the formation of large, well-ordered crystalline regions. Instead, the polymer solidifies in an amorphous state, where molecular chains are randomly arranged without long-range order ^49^.

In this work, we introduced a thermal quenching step in the casting process of the polymer by rapidly bringing it to room temperature through exposure to air rather than letting the casted material slowly cool down in the oven (Fig. S1a).

The effectiveness of the quenching was then evaluated by using DSC to compare the starting granules and the polymer after casting and quenching (Fig. S1b).

The presence of two distinct peaks in the DSC trace of the granules, with one peak around 65°C and the other around 170°C, indicates different thermal transitions occurring within the polymer. The peak at ~65°C likely corresponds to the glass transition temperature of the polymer, while the one at ~170°C is typically associated with the melting temperature of the crystalline regions within the polymer. Crystalline polymers have ordered molecular arrangements in certain regions, and at the melting temperature, these regions lose their order and transition into a disordered, liquid-like state ^50^. In comparison, the crystalline peak at around 170°C is suppressed in the DSC curve of the cast and quenched PDLLA because there are no well-defined crystalline regions to melt during heating, confirming the efficacy of this casting protocol.

The obtained amorphous PDLLA bulk was then used to fabricate preforms with a diameter of 10 mm by mechanical machining, which were then scaled in diameter down to sub-millimeter optical fibers by a standard thermal drawing process (Figure 1a). We reduced the initial preform diameter by ratios between ~20 and ~70 times, resulting in several tens of meters of fiber obtained in a single drawing, with diameters ranging from 150 to 510 μm (Figure 1b (I-III)). The transmittance of the drawn fibers was assessed for different fiber diameters by using a broadband supercontinuum laser as the light source (Figure 1c). The measured spectra show a broad transmission band ranging from the visible (red light) to ~1650 nm, with two marked absorption bands, attributed to the second overtone of C-H stretches (1110-1230 nm) and the first overtone of O-H bonds (1330-1430 nm) ^51^. In particular, one of the spectral regions with the highest transmittance overlaps with the near-infrared (NIR) tissue transparency spectral region (~650-900 nm) ^52^.

**Figure 1:**
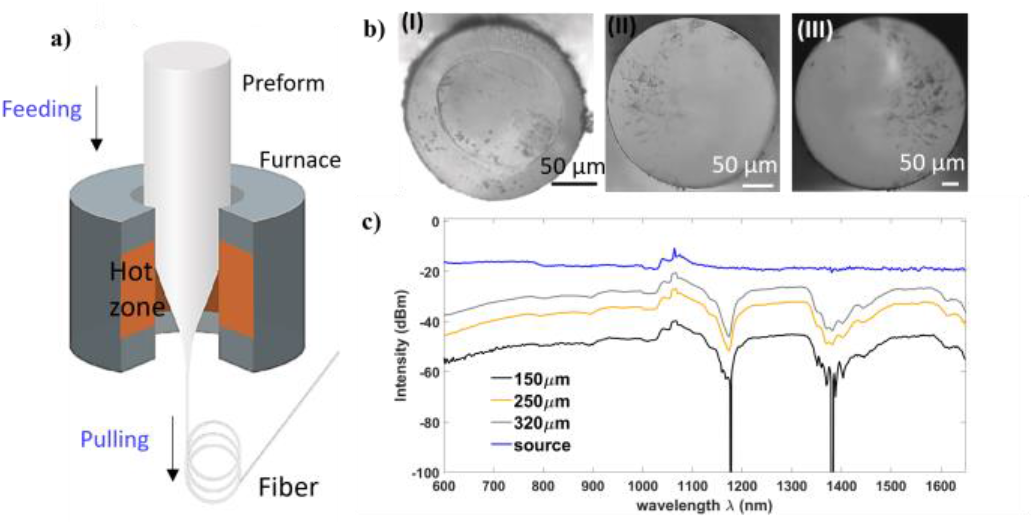
**(a)** Schematic of the thermal drawing process used to fabricate the biodegradable optical fibers. **(b)** cross-sectional microscope images of drawn PLA fibers with diameters of (I) 150, (II) 250 and (III) 320 μm, respectively. **(c)** transmission spectra of 5-cm-long fiber pieces of different diameters; in comparison with the spectrum of the supercontinuum laser used as source.

A direct comparison between the spectra in Figure 1c reveals a trend of increased transmission with increased fiber diameter, attributed to the variations in mode coupling. Smaller diameter fibers typically support fewer propagation modes, while larger diameter fibers support a very large number of modes, leading to increased mode coupling ^53^.

### Optical fiber degradation in PBS

The first validation of the biodegradability of the fabricated optical fibers was performed by measuring their optical and morphological properties after immersion in PBS for up to 21 days.

During the degradation process, especially since the fiber is unstructured (no separate cladding material), the increase in surface porosity and fluid permeation within the fiber is expected to increase light scattering and subsequently negatively impact light propagation through the fiber. We experimentally measured this phenomenon by comparing the transmission spectra of an as-drawn fiber and a fiber soaked in PBS for 14 days (Figure 2a, and Figure S1c). As expected, we observed a marked decrease in transmittance in the sample that had been immersed in the solution.

**Figure 2:**
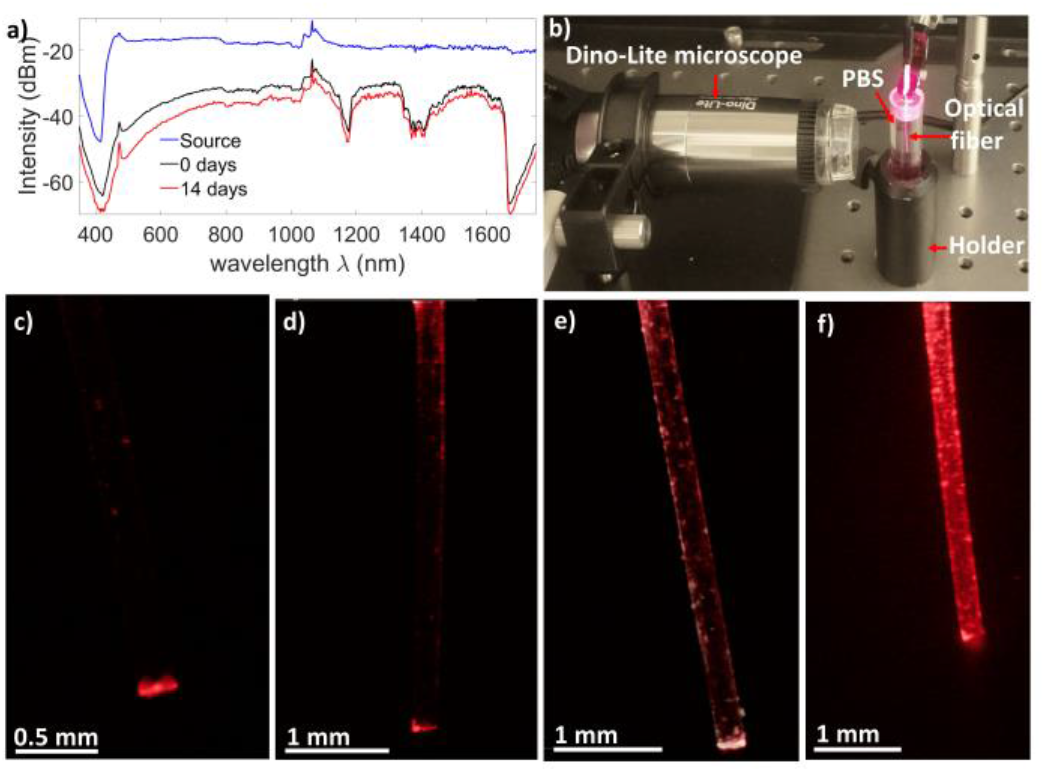
**(a)** In black and red, comparison between the transmission spectra of 5-cm-long pieces of fibers before and after 14 days of soaking in PBS; in blue, the spectrum of the supercontinuum laser used as source. **(b)** Changes in light scattering from the side portions of the immersed optical fiber in PBS, recorded using a Dino-Lite optical micro-scope, demonstrating the effects of degradation. **(c-f)** Digital microscope images of a fiber guiding red light while immersed in PBS for 0, 7, 14 and 21 days, respectively.

Moreover, the increase in scattering during degradation was confirmed by using a digital microscope to monitor the evolution of the side-scattered light intensity over 21 days of immersion in PBS (Figure 2b). As visible from Figure 2c, at the moment of immersion, most of the light could be observed at the fiber’s output facet. After 7 days a significant amount of scattering points was visible on the fiber’s outer surface (Figure 2d). The bulk degradation further increased in the following two weeks (Figure 2e), and by day 21 most of the light recorded by the microscope was scattered from the fiber’s sides (Figure 2f).

The faster degradation between days 14 and 21 could be explained by the fact that PDLLA degradation involves hydrolysis, a process where water molecules break the ester bonds in the polymer chain, leading to a loss of mechanical integrity ^54 55^. As the polymer degrades, it may become more porous and absorb more water, enhancing the availability of water molecules for hydrolysis reactions. This increased water absorption in the third week could contribute to a faster degradation rate ^56^. Moreover, the initial heterogeneous degradation of PDLLA, which involves several competing mechanisms such as hydrolysis ^57^, chain scission ^58^, and other chemical and physical reactions ^59^, may result in changes to the surface area of the fiber, potentially influencing the rate of degradation. Changes in surface area might become more significant as the degradation progresses, contributing to an accelerated degradation in the later stages of the process ^60^.

### In vivo evaluation of degradability

Following the promising in vitro results, we proceeded to evaluate the behavior of our biodegradable fibers in living neural tissue. To do so, fibers with different diameters of 510, 290, and 210 μm were surgically implanted in the LP region of the brain of freely behaving mice for periods of 2 and 4 months (Figure 3a(I)).

**Figure 3:**
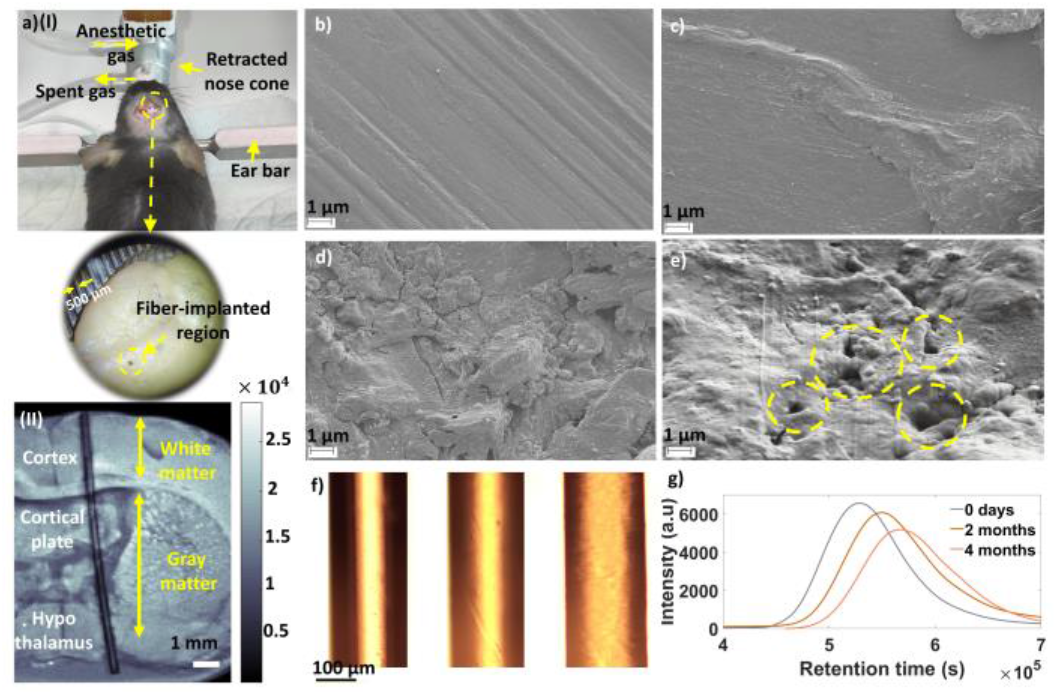
**(a)** (I) PLA optical fibers were implanted in the Lateral posterior nucleus of the thalamus (LP) region of the mice brains (n=2). b-d) SEM image of the morphology of the fiber surface before implantation **(b)** compared with the morphology after 2 and 4 months of implantation **(c)** and **(d)**, respectively). **(e)** porous structure of the implanted PLA fiber in the gray matter of the mouse brain after 2 months. **(f)** Polarized light optical image of the fiber at different stages of degradation; from left to right: pristine, after 2 months of implantation and after 4 months of implantation. **(g)** Comparison between the molecular weights of the implanted fibers after 0 days, 2 months, and 4 months, by GPC mass spectrometry.

The thickness of the white matter in the targeted region was approximately 2.56 mm. Therefore, we implanted 7 mm long optical fibers to gain access to both the white matter and gray matter, allowing us to compare the degradation between the two. After the implantation periods, the animals were transcardially perfused, the fibers were explanted, and their brains were dissected and sliced in coronal sections.

In order to visualize the size of the implant in relation to different brain structures, we re-implanted the optical fiber in the brain slice and used a custom-made mid-infrared photoacoustic microscope (MIR-PAM) (Figure 3a(II)) to image it. We chose MIR-PAM as an imaging technique drawing from previous findings from FTIR spectroscopy studies on PDLLA ^61^. These studies consistently demonstrated heightened absorptions in the MIR region, attributed to key functional groups within PDLLA’s molecular structure, such as C-H stretching bonds, carboxyl groups, CH3 deformation modes, and ester groups. By using a laser with a wavelength of 3.4 μm, which corresponds to the absorption peak of C-H stretching bonds in PDLLA at 2941 cm-1, we intended to take advantage of the fact that PDLLA absorbs light strongly at this specific wavelength. This absorption allows for enhanced signal-to-noise ratio and specificity when visualizing the PDLLA implant using MIR-PAM.

In previous research, evaluating and tracking the degradation of implanted biodegradable polymers often relied on labeling the polymers with carbon-14 isotopes (C-14) ^62–64^. However, this approach has several drawbacks. Primary among these are concerns about radioactivity, which requires strict adherence to safety protocols and regulations. Additionally, measurements can be inaccurate due to interference from background radiation, and there are limitations in the availability of C-14 isotopes ^65^.

In our current study, we employed alternative methods to assess the degradation profile of the implanted polymer optical device: SEM imaging, optical microscopy, and molecular weight measurements.

Firstly, after removing the fibers from the brain tissue, we performed SEM imaging of the fibers’ surfaces (Figure 3b-d). A direct comparison between the as-drawn fiber and the fiber implanted for 2 months shows a visible degradation of the fiber’s surface, which turns into a complete erosion after 4 months. Interestingly, as shown in Figure 3e and S2, this effect appears to be more marked in the part of the fiber that had been in contact with grey matter region of the brain (Figure S2(I)) when compared to the parts in contact with white matter (Figure S2(II)). This outcome can be attributed to the different enzymatic environments in the two kinds of brain regions. Enzymatic degradation of PDLLA may involve enzymes such as esterases ^66^. Recent studies showed that the concentration of these enzymes, which catalyze the hydrolysis of ester bonds ^67^, was higher in gray matter compared to white matter ^68^. This is due to the greater metabolic activity and neuronal density in gray matter, which may require greater esterase activity for the breakdown of acetylcholine and other ester-containing molecules ^69^. Hence, the concentration of esterases, including acetylcholinesterase and other esterases that can degrade PLA, is generally higher in gray matter compared to white matter in the brain. Therefore, the difference in the porous structure on the fiber, as depicted in Figure S2(III) and Figure 3e, indicates that the degradation of the polymer occurs more rapidly in the gray matter compared to the white matter after 2 months of implantation.

Degradation of the polymer’s surface, as well as increased porosity due to the volumetric degradation within the fibers and formation of H2O and CO2 bubbles as a result of secondary hydrolysis ^70^, are expected to increase the overall scattering coefficient. This effect can be visually observed by optical transmission microscopy with polarized light, since multiple scattering can randomize polarization, resulting in a decrease in the detected light intensity when the polarizer and analyzer are aligned ^71^.

Thus, the progressive reduction of the maximum intensity in the optical microscope images presented in figure 3f further confirms the in vivo degradation of the optical fiber (Figure 3f).

In addition to the increase in porosity, as the degradation process of the implanted polymer initiates from the surface of the optical fibers, microscopic portions of material detach from the bulk of the device, resulting in morphological changes. Figure S3a(I) illustrates the smooth surface of the non-degraded PDLLA optical fiber, contrasting with the degraded surface of the implanted PDLLA optical fiber after 2 months in the brain of the animal (Figure S3a(II)).

Finally, the degradation of the fiber on a molecular level was estimated by gel permeation chromatography (GPC), a reliable technique commonly used to measure the molecular weight distribution of polymers ^72,73^. Compared to the pristine polymer, we observed higher average retention times for samples degraded in the brain tissue, corresponding to lower molecular weights, as well as broadened distributions of the retention times (Figure 3g). These results are compatible with the non-uniform shortening of the polymeric chains caused by the breaking of bonds between chemical groups by either hydrolysis or enzymatic reactions, which are the main mechanisms underlying the degradation of PDLLA in living tissue on a molecular level

## Conclusion and future prospect

In this study, we fabricated an amorphous PDLLA optical fiber by combining casting and quenching of granules with thermal drawing. The light transmission profile of the fabricated polymer fiber shows high transmission in the SW-NIR region. This paves the way for applications within this wavelength range, which is known to allow for greater tissue penetration when compared to visible light due to reduced light scattering. An in vitro optical investigation of the degradation behavior of the fiber revealed an increase in scattering during light propagation, compatible with increased porosity and damage to the surface caused by hydrolysis of the polymeric chains. Morphological analysis of fibers implanted in the brains of freely behaving mice for up to 4 months was used to verify in vivo degradation and highlighted a dependence of the dissolution rate on the implantation region due to different enzymatic processes in gray and white matter. Finally, the in vivo degradation was confirmed by gel permeation chromatography, which showed a decrease in the PDLLA’s molecular weight over the course of implantation.

The biodegradable optical fiber presented in this study has the potential to be used for multifunctional and diverse applications in biomedical experiments, including optogenetic stimulation, fluorescence photometry, phototherapy, and laser surgery. Overall, the use of the PDLLA POFs in implantable devices holds great promise for the development of advanced implantable technologies that can improve patient outcomes and quality of life.

## Acknowledgment

The authors acknowledge financial support from the Lundbeck Foundation (Multi-BRAIN, R276-2018-869 and R380-2021-1171) and the Villum Fonden (36063). We would like to express our gratitude to Lotte Nielsen (Laboratory Technician, Department of Health Technology), Berit Herstrøm (Process Specialist, M.Sc., National Centre for Nano Fabrication and Characterization Process Engineering). Also, Richard Crane and Jakob Janting from DTU Electro assisted with the DSC measurements.

## Materials and methods

### Preform fabrication and thermal drawing process

200 g of PDLLA granulate (nominal granule size: 3-5 mm, melt flow rate: 24, extrusion grade, D-content of 2 %, Goodfellow GmbH) were dehydrated under vacuum at a temperature of 50°C for 12 hours. Subsequently, the granules were cast under vacuum in an aluminum mold at 230°C for 24 hours. To achieve further dehydration, the vacuum oven’s temperature was incrementally increased in two stages, commencing from room temperature and progressing to 100°C for 12 hours, followed by an increase to 230°C. The molten PDLLA material was finally air-quenched within a desiccator, maintaining a vacuum at room temperature for 48 hours. In all the above procedures, the role of the vacuum environment was to prevent the formation of bubbles within the cast material. The consolidated PDLLA was then mechanically machined into rods measuring 10 mm in diameter and 10 cm in length. These rods were further dehydrated in a vacuum oven at 60°C for 72 hours before the fiber fabrication, which was achieved by the thermal drawing process of the rods in a custom-made draw tower, as outlined in our previous studies ^18,74^. During the drawing, the temperature gradually increased from room temperature to 185°C, at a rate of 1°C/min. The diameter of the fiber was controlled by regulating the capstan speed and feeding speed, following the provided equation:

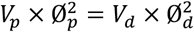

(Vp: feeding speed; Vd: drawing speed; Øp: preform diameter; Ød: fiber diameter). The diameter of the fiber was accurately measured during drawing by a laser detector with a resolution of <0.1 μm. Using this process, we fabricated fibers having different diameters between 150 and 510 μm.

### Differential scanning calorimetry (DSC)

Thermograms for the PDLLA granules as bought and for the polymer bulk obtained by casting and quenching were measured with a Q 2000 DSC (TA Instruments, USA). Prior to testing, the samples (10 mg each) were subjected to vacuum drying for a duration of 48 hours. The measurements were performed in the −10 to 200°C temperature range, with a heating rate of 10°C/min and a cooling rate of 50°C/min. The measurement was repeated three times for each sample.

### Broadband transmission measurements

The optical transmission of the PDLLA fiber was characterized by a broadband supercontinuum source (NKT SuperK Extreme) in the range of 400 - 1700 nm. The light from the supercontinuum source was reduced to avoid thermal damage to 25 mW (500KHz repetition rate), and butt-coupled to PDLLA fibers with a length of 5 cm and different diameters using a silica patch cable with a core diameter of 105 μm (Thorlabs M61L01). The spectra at the output of the PDLLA fibers were recorded by an Ando AQ-6315A optical spectrum analyzer. The same setup was also used to evaluate the transmission properties of the PDLLA optical fiber (d=230μm, L=5cm) before and after in vitro degradation in PBS for 2 weeks.

### Evaluation of side scattering in PBS

The light scattered from the side of the optical fiber, the intensity of which is correlated to the material’s degradation, was imaged continuously using a digital microscope (Dinolite AM7915MZT) over 3 weeks of submersion of the fiber in PBS (pH 7.4). The sample used for this measurement is a 230 μm diameter PDLLA fiber (L=5cm) connectorized with a ceramic ferrule (Thorlabs CFLC230) and coupled to a LED source (Thorlabs M660L4).

### Microscopy

Optical images of the cross-sections of fibers with different diameters were acquired using a Zeiss Axioscan A1 microscope after polishing the end facets of short pieces of fiber on a sequence of different lapping films (30, 9, 6, 3, 1.5, 1, and 0.5 μm grades, respectively). Side images of the fibers before and after degradation in PBS were obtained using the same microscope. Images of the fibers implanted in the brain were instead acquired using a Bresser Science ADL 601 P optical microscope equipped with a polarizer and analyzer for evaluation of the material’s scattering. SEM images of the fibers’ surface were performed using a ZEISS EVO modular SEM platform (voltage: 1.35 KV, magnification: 3.00 KX, pixel size: 97.72 nm).

### Gel permeation chromatography (GPC)

A gel permeation chromatograph (HLC-8320, TOSOH, Japan) was employed to compare the molecular weight of PDLLA before and after in vivo degradation. The samples used (N=3 for each case) were solutions prepared by mixing 2.5 mg of PDLLA in 1 ml of chloroform (Shimadzu, Germany). The mobile phase was maintained at a flow rate of 0.6 mL/min, the column temperature was set to 40°C, and each injection consisted of a 10 μL volume.

### Surgical procedures

We employed a surgical protocol similar to the one outlined in our previous study ^21^. All in vivo experiments were performed on wild-type mice (female, age: 1 month, ~30g), using aseptic surgical conditions. A total of 2 animals, each representing an experimental unit, were employed in this study, corresponding to the two implantation periods to be compared (2 and 4 months). The implantation of the biodegradable fibers was performed under isoflurane gas (1.5-2%) anesthesia (0.2 ml/min oxygen pressure, and 0.3 ml/min air pressure). After head shaving, the animals were placed in a stereotaxic frame, where their head was fixed by ear bars. An incision was made on the head after sterilizing the skin with ethanol (70%) and injecting lidocaine in situ (0.01 ml) for local anesthesia. A hemostat was used to keep the incision open, and the bleeding was controlled by using hydrogen peroxide, cautery, and saline ^75^. A drill was used to make a hole in the skull, and the dura was removed using a needle under a surgical microscope to access the region of interest in the brain. The PDLLA polymer fibers (7 mm in length, 210 μm, 290 μm and 510 μm in diameter) were implanted in the LP region of the brain, using the robotic arm of the stereotaxic frame. The hole was sealed after implantation with dental cement and cyanoacrylate adhesive, and the incision was sutured. Here, we considered the optical fibers that were not implanted, but were of the same size as the implanted optical fibers, as the control groups.

### Post-surgical care

Immediately after surgery, the mice were kept on a heating pad (37°C) and injected with 0.003 ml of Baytril, 0.003 ml of Carprofen, 0.0048 ml Bupad, and 2-3 ml saline. The mice were then placed in separate cages post-surgery to prevent interference from cage mates. The cages had proper ventilation, gentle lighting, and adhered to a light-dark cycle. Soft and comfortable bedding materials were placed in the cages. A relative humidity level between 45% and 65% was maintained. The mouse rooms and cages were kept within a temperature range of 20-24°C. The mice were administered a mixture of Buprenorphine (0.2 mg) and Nutella (1g) (B+N) at a dosage of 0.4 mg/kg, 8 hours after surgery. Additionally, Carprofen (0.003 ml) was injected 12 hours after the initial feeding, followed by another B+N feeding. This process was repeated at 12-hour intervals for a total of 4 days post-surgery.

### Brain extraction and slicing

At the end of the implantation periods (2 to 4 months), the animals were deeply anesthetized by intraperitoneal injection of a mixture of ketamine (80 mg/kg) and xylazine (10 mg/kg). The mice were then transcardially perfused with 10% formalin solution in PBS. For the perfusion procedure, the chest cavity was opened, and saline was injected into the left ventricle, using a 30-gauge needle. The right ventricle was cut immediately after injection. The perfusion was continued with formalin solution for another 10 minutes. Afterward, the animals were decapitated, and the brains were extracted by cutting the cranium. The extracted brains were first kept in formalin solution (10%) for 24 hours, followed by transfer to PBS (pH= 7.4) at 4°C for storage. The brains were finally sliced into thin sections (thicknesses of 0.5 and 1 mm) using a Zivic brain slicer matrix.

All the procedures concerning the housing of rats, habituation in the animal facility, surgery, post-surgical care, and euthanasia were standardized to minimize variability, approved by the National Veterinary and Food Administration (permission number for the animal research: P22-477), and in compliance with the European Union Directive 2010/63/EU.

### Photoacoustic imaging (PAI)

PA images of the PDLLA optical fiber (initial diameter of 290 μm) embedded in a brain slice (500 μm thickness) after 2 months of in vivo implantation were acquired using an inhouse transmission mode MIR-PAM setup. A 3.4 μm wavelength laser focused at the bottom of the sample, which was submerged in water, was used as an excitation source. The resulting ultrasounds were collected by an immersive focused transducer (20 MHz center frequency, 8 mm focal length, 10 mm diameter) on the top. The acquired photoacoustic signal underwent filtering through two analog filters (Mini-circuits) – a 1 MHz low-pass filter and a 27 MHz high-pass filter. After filtering, the signal was amplified using dedicated amplifiers and subsequently fed into a fast digitizer (M4i.4421-x8, Spectrum Instrumentation), operating at a sampling rate of 250 MS/s (mega-samples per second), for data processing. To ensure precise scanning of the sample, we employed a high-resolution XY stage (8MTF-75LS05, Standa) with movement control provided by a stepper and DC motor controller (8SMC5-USB, Standa). The images were acquired with a scanning step size of approximately 50 μm.

## Data analysis

All data analysis and visualization were performed using the MATLAB suite. No experimental unit was excluded when analyzing the results of in vivo experiments.

## Supplementary Information

**Figure S1:**
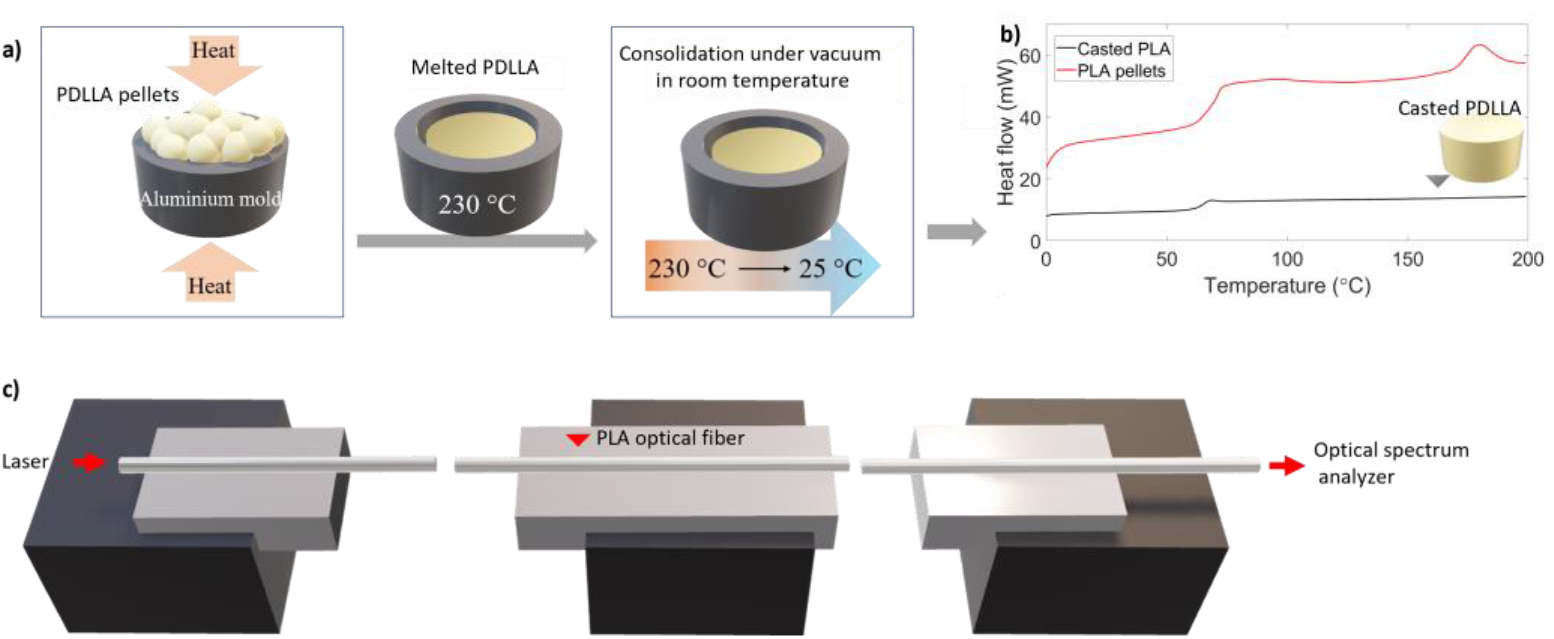
**(a)** PLA granulate casting in the vacuum oven at 230°C for 24h, followed by air quenching to 25°C. **(b)** DSC results of the PDLLA pellets in comparison with the quenched casted PDLLA **(c)** Optical system for light transmission measurement.

**Figure S2:**
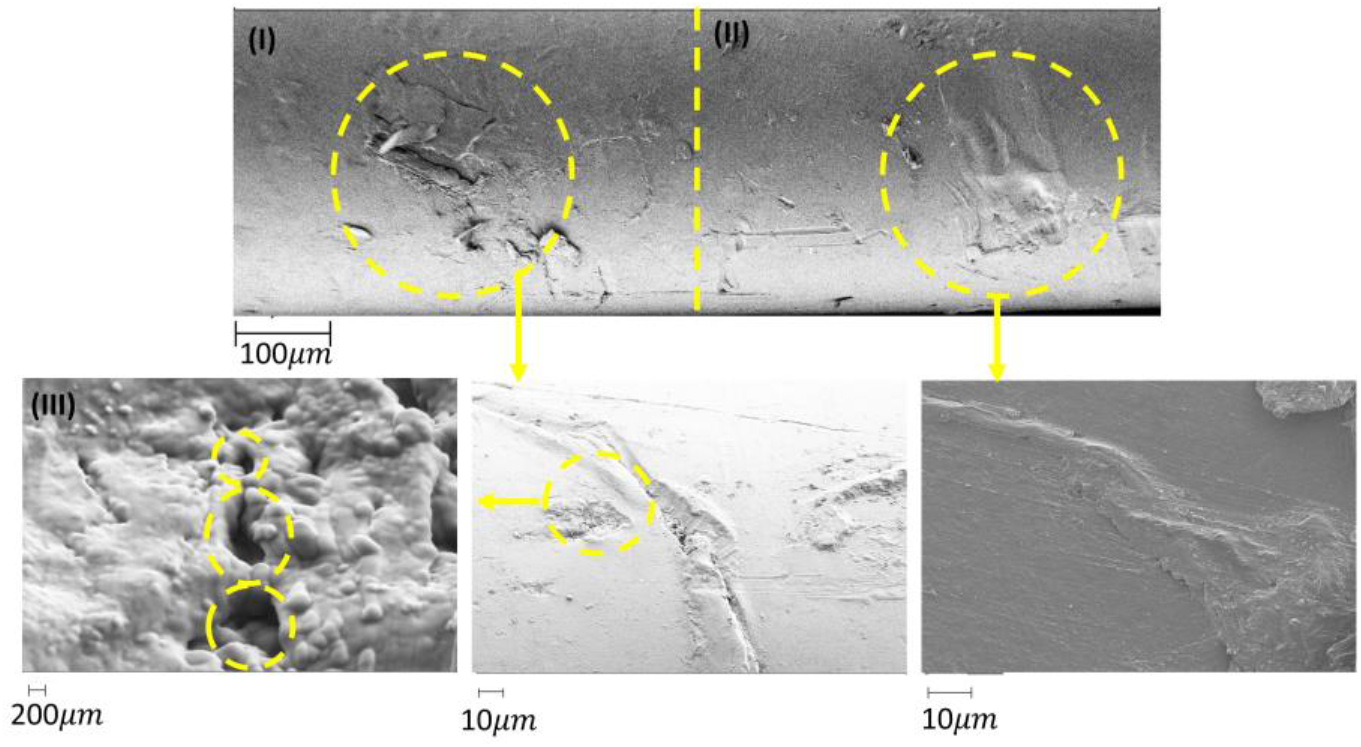
Comparison between the degradation of the polymer fiber in **(I)** white matter, and **(II)** gray matter. **(III)** porous structure appeared on the PLA surface after 2 months of in vivo implantation in the gray matter of the brain.

**Figure S3:**
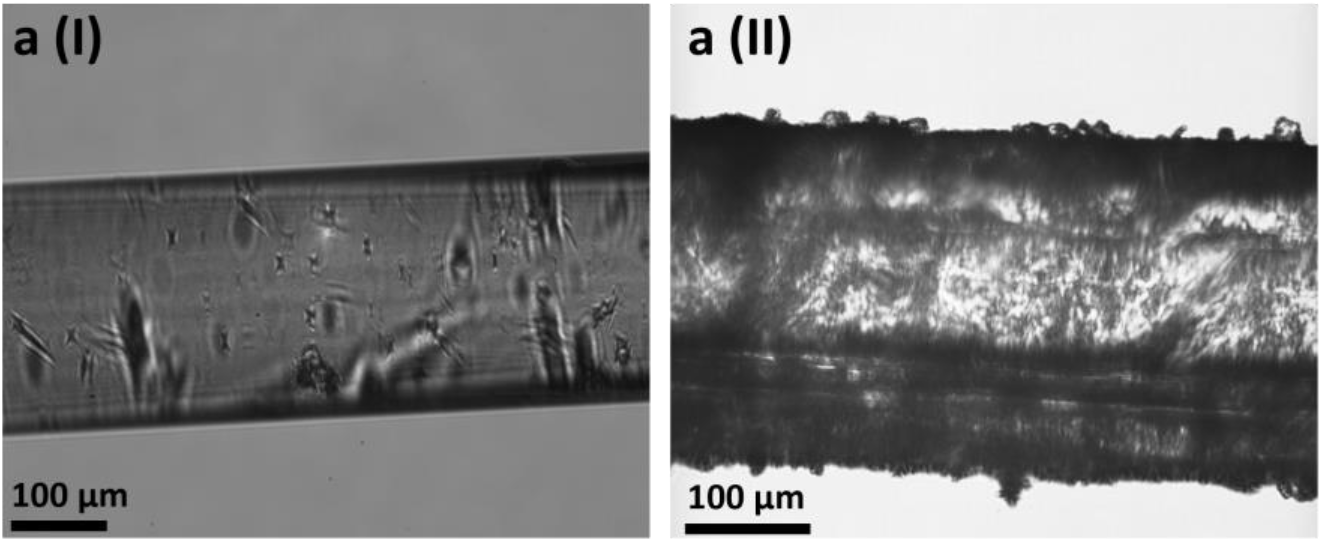
**(a(I))** Morphological changes of the surface of the non-implanted optical fiber in comparison with that for **(a(II))** the 2-month-implanted optical fiber in the brain.

